# An amplicon-based nanopore sequencing workflow for rapid tracking of avian influenza outbreaks, France, 2020-2022

**DOI:** 10.1101/2023.05.15.538689

**Authors:** Guillaume Croville, Mathilda Walch, Laetitia Lèbre, Sonia Silva, Fabien Filaire, Jean-Luc Guérin

## Abstract

During the recent avian influenza epizootics that occurred in France in 2020/21 and 2021/22, the virus was so contagiousness that it was impossible to control its spread between farms. The preventive slaughter of millions of birds consequently was the only solution available. In an effort to better understand the spread of avian influenza viruses (AIVs) in a rapid and innovative manner, we established an amplicon-based MinION sequencing workflow for the rapid genetic typing of circulating AIV strains.

An amplicon-based MinION sequencing workflow based on a set of PCR primers targeting primarily the hemagglutinin gene but also the entire influenza virus genome was developed. Thirty field samples from H5 HPAIV outbreaks in France, including environmental samples, were sequenced using the MinION MK1C. A real-time alignment of the sequences with MinKNOW software allowed the sequencing run to be stopped as soon as enough data were generated. The consensus sequences were then generated and a phylogenetic analysis was conducted to establish links between the outbreaks.

The whole sequence of the hemagglutinin gene was obtained for the 30 clinical samples of H5Nx HPAIV belonging to clade 2.3.4.4b. The consensus sequences comparison and the phylogenetic analysis demonstrated links between some outbreaks.

While several studies have shown the advantages of MinION for avian influenza virus sequencing, this workflow has been applied exclusively to clinical field samples, without any amplification step on cell cultures or embryonated eggs. As this type of testing pipeline requires only a short amount of time to link outbreaks or demonstrate a new introduction, it could be applied to the real-time management of viral epizootics.

## Introduction

Avian influenza is a highly contagious infectious disease caused by Influenza A viruses belonging to the *Orthomyxoviridae* family ^1^. Aquatic wild birds represent the natural reservoir of the virus, contributing to its spread and the generation of occasional epidemics characterized by the deaths of both wild and domestic birds, generating significant economic losses ^2,3^.

The highly pathogenic avian influenza viruses (HPAIVs) responsible for the 2020/21 and 2021/22 European epizootics belong to the H5N8 and H5N1 subtypes, both belonging to the A/Goose/Guangdong/1/1996 (GSGd) lineage. The HPAIVs actually circulating in Europe descended from the H5N8 variant (GsGd lineage of clade 2.3.4.4b) that emerged in Europe in 2014 after an intercontinental spread which started in Southeast Asia in early 2014 ^4–8^.

As of the 1^st^ of March 2023, the 2020/21 epizootics led to 3,555 reported HPAI detections and affected around 22,400,000 poultry birds in 28 European countries ^9^. In 2021/22, the joint EFSA, ECDC and EU reference laboratory report listed a total of 2,467 outbreaks in poultry, 48 million birds culled in the affected establishments, 187 detections in captive birds, and 3,573 HPAI events in wild birds, affecting 37 European countries ^10^. The 2021/22 outbreak is considered the largest and most devastating HPAI outbreak ever to occur in Europe and, more recently, in the United States, where 46 American states were affected, resulting in 49 million dead birds (death as a result of AIV infection or culling) ^11^.

Because avian influenza viruses are highly contagious, and given the potential risk of long-distance spread of HPAI viruses from infected barns, it is crucial to conduct real-time genetic investigations to track circulating AIV strains. This real-time genetic monitoring of circulating strains aims to identify possible links between outbreaks in order to ultimately break the chain of infection.

Several recent studies focused on pipelines based on next-generation sequencing (NGS) to detect influenza viruses [12–19]. However, none of these studies used nanopore sequencing of clinical samples or open source bioinformatics algorithms to identify epidemiological links between spontaneous HPAIV outbreaks.

This work aims to combine a multiplex PCR amplification of H5 avian influenza viruses with nanopore sequencing to evaluate the pathotypes and clades of circulating viruses, and identify the genetic links between 11 outbreaks that occurred in France between 2020 and 2022.

## Results

### *In silico* validation of PCR-based enrichment

An *in-silico* analysis consisting in the alignment of 20 PCR primers (Table 1) selected in the literature with a panel of H5 hemagglutinin was first performed for the constitution of the multiplex PCR panel. The selection of the PCR primers was based on their hybridization locations across the HA segment to guarantee a whole or partial gene amplification including the cleavage site. We also included MBTuni-12 and MBTuni-13 universal primers ^12^ for the full amplification of any AIV subtype. The amplicon sizes produced by the different combinations of PCR primers are available in Table 2.

**Table 1.**
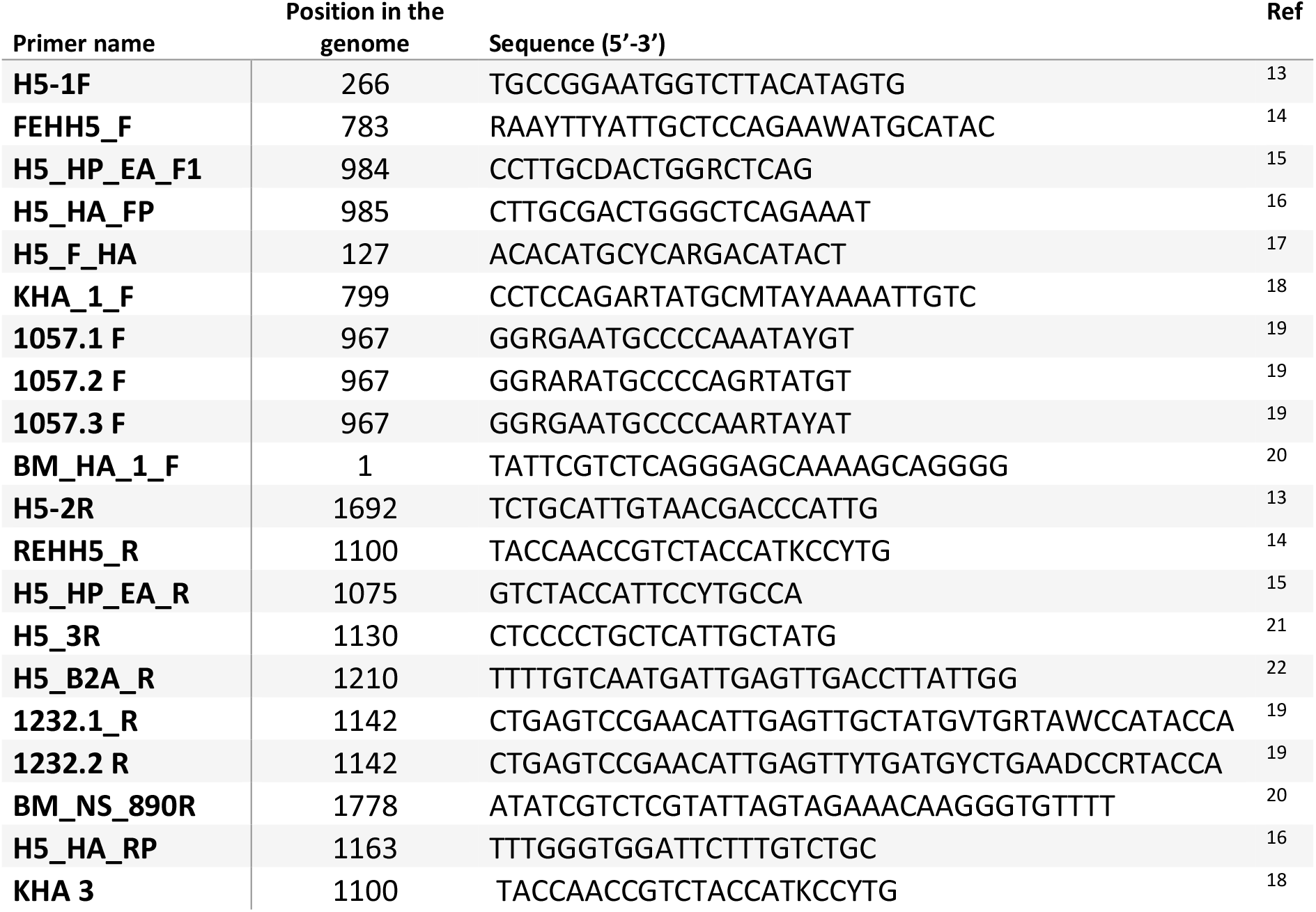
Names, positions, sequences and references of the PCR primers used in this study.

**Table2.** Hemagglutinin amplicons sizes (bp) according to primer combination.

| Rev primer name →<br>For primer name ↓ | H5-2R | REHH5 R | H5 HP EA R | H5 3R | H5 B2A R | 1232.1 R | 1232.2 R | BM NS<br>890R | HA RP | KHA 3 |
| --- | --- | --- | --- | --- | --- | --- | --- | --- | --- | --- |
| **1057.1 F** | 1426 | 834 | 809 | 864 | 944 | 876 | 876 | 1512 | 897 | 834 |
| **H5 HP EA F1** | 909 | 317 | 292 | 347 | 427 | 359 | 359 | 995 | 380 | 317 |
| **KHA 1 F** | 708 | 116 | 91 | 146 | 226 | 158 | 158 | 794 | 179 | 116 |
| **FEHH5 F** | 707 | 115 | 90 | 145 | 225 | 157 | 157 | 793 | 178 | 115 |
| **H5 F HA** | 1565 | 973 | 948 | 1003 | 1083 | 1015 | 1015 | 1651 | 1036 | 973 |
| **1057.3 F** | 893 | 301 | 276 | 331 | 411 | 343 | 343 | 979 | 364 | 301 |
| **H5 HA FP** | 725 | 133 | 108 | 163 | 243 | 175 | 175 | 811 | 196 | 133 |
| **H5-1F** | 725 | 133 | 108 | 163 | 243 | 175 | 175 | 811 | 196 | 133 |
| **1057.2 F** | 725 | 133 | 108 | 163 | 243 | 175 | 175 | 811 | 196 | 133 |
| **BM HA 1 F** | 1692 | 1100 | 1075 | 1130 | 1210 | 1142 | 1142 | 1778 | 1163 | 1100 |

### Validation of the multiplex-PCR sequencing on different AIV subtypes

The evaluation of the PCR panel was then performed on several AIV subtypes of highly pathogenic (H5N8) and low pathogenicity avian influenza viruses (H5N3, H6N1, H7N1 and H9N2). These isolates were submitted to our nanopore protocol: RNA extraction, PCR amplification and ONT sequencing. These assays resulted in a complete coverage of the HA gene and a near-complete to whole sequencing of the seven other RNA segments (Table 3).

**Table 3.** Size (bp) of the consensus sequence obtained by nanopore sequencing on different AIV subtypes.

| AIV subtype | NCBI<br>taxon ID | CT value<br>(M gene) | PB2 | PB1 | PA | HA | NP | NA | M | NS |
| --- | --- | --- | --- | --- | --- | --- | --- | --- | --- | --- |
| **H5N3** | 437394 | 32 | 290 | 1109 | 498 | 1711 | - | 263 | 943 | 845 |
| **H6N1** | 1096539 | 32,41 | 457 | 1338 | 2158 | 1711 | 312 | 1364 | 989 | 845 |
| **H7N1** | 437402 | 31,84 | 2121 | 2110 | 2158 | 1693 | 1501 | 1354 | 989 | 845 |
| **H9N2** | OQ179925<br>(access. number) | 30,23 | 638 | 1101 | 998 | 161 | 1501 | 291 | 989 | 845 |

The resulting PCR products obtained from two H5N8 sample were loaded on an agarose gel to demonstrate the amplification of the HA in several parts, as well as the whole genome amplification with the universal primers (Fig. 1).

**Figure 1.**
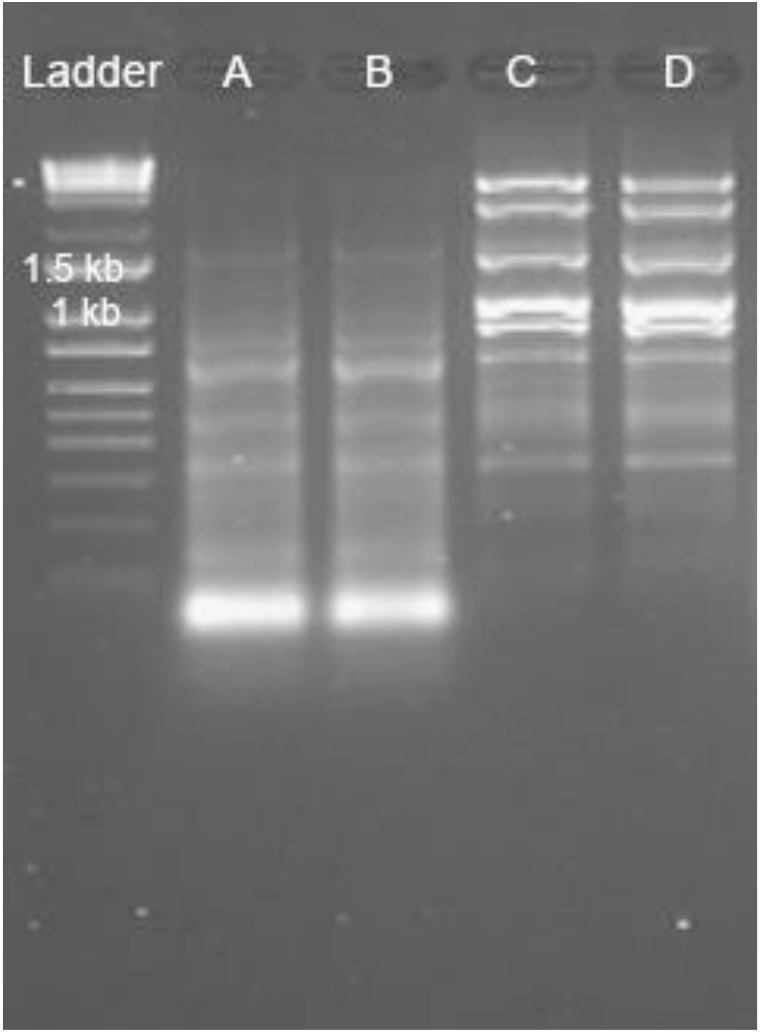
Amplicons profile on a 1% agarose gel. The resulting PCR products obtained from two H5N8 samples were loaded on an agarose gel. The DNA bands show the amplification of the HA with the HA primers pool in several parts (A, B), as well as the whole genome amplification with the universal primers (C,D). This image represents a cropped gel.

### Application of the multiplex-PCR sequencing pipeline on two sets of field samples originating from 11 H5N8 and 19 H5N1 HPAIV outbreaks sampled in 2020-2021 and 2021-2022, respectively

Two sets of field samples were included in this study, both originating from French outbreaks: 11 from December 2020 to February 2021, and 19 from late November 2021 to February 2022 (Supplementary File S1). The Ct values ranged between 13.03 and 28.5, which is representative of high and low viral loads, respectively. These field samples included mostly tracheal swabs, but also feather pulp, cloacal swabs and dust wipes, to cover the different types of samples available for AIV surveillance.

Each sample was submitted to HA and whole-genome amplification using the multiplex PCR pipeline described previously. The sequencing yields varied from 2.34 to 474.13 Mb per sample, while the N50 values were comprised between 469 and 1 818 bp. The sequencing metrics such as read length, N50 and number of reads for each sample are available in Supplementary File S1. An example of cumulative output (bases) is shown in Fig. 2A, corresponding to a run where four samples were multiplexed on a flongle flowcell. Fig. 2B and 2.C show a read length histogram and the bases count per barcode, respectively. Finally, Fig. 2D shows the coverage and sequencing depth obtained for one sample; the 30 figures corresponding to the 30 sequenced samples are available in Supplementary File S2.

**Figure 2.**
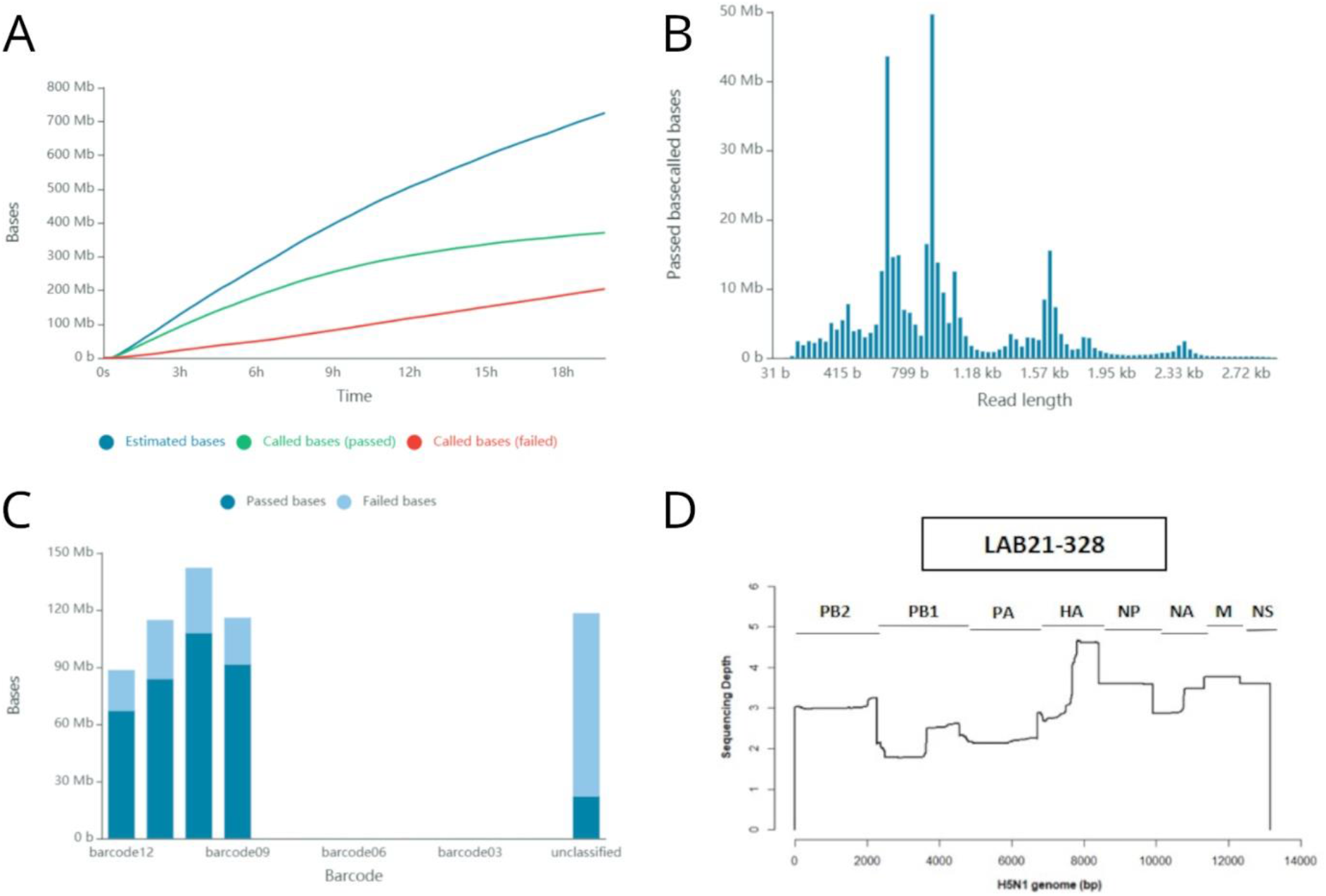
Sequencing metrics and genome coverage. Fig. 2A shows the cumulative output of bases produced as a function of time (hours). Fig. 2B shows a read length histogram with several peaks corresponding to the size of the majority amplicons. Fig. 2C shows the bases count per barcode in a sequencing run where 4 samples were multiplexed. Fig. 2D shows the coverage and sequencing depth all over the whole genome obtained for one sample.

The consensus sequences were obtained for each hemagglutinin, using iVar, and were deposited in the Genbank database. HA accession numbers are listed in Supplementary File S1, and the complete list of accession numbers is available in Supplementary File S3.

### Phylogeny

Two time-scaled phylogenetic trees (Supplementary files S4 and S5) were built in order to determine whether the outbreaks were related to each other (e.g. farm network) or were due to new viral introductions (e.g. an unnoticed outbreak, introduction from wild birds). Supplementary file S4 shows the relationship between nine outbreaks (blue taxa) in southwest France (Gers, Hautes-Pyrénées, Landes) between 6 December 2020 and 20 February 2021. In addition, at a distance of about 300 km (in Vendée and Deux-Sèvres), two other outbreaks (green taxa) were sampled on 13 December 2020. The tree in Supplementary files S4 shows a common recent ancestor to the 11 French sequences, but it is clear that the nine outbreaks in the southwest are not directly related to the outbreaks in Vendée and Deux-Sèvres. Supplementary file S5 shows 19 outbreaks clustering in six different groups. A set of 11 sequences (pink taxa) from southwest France sampled over a 4-week period (18 December 2021 to 19 January 2022) illustrates the circulation of one viral strain within a restricted area, suggesting a link between these farms that could be the source of the virus circulation. In addition, in the same area and approximately at the same time, we analyzed six outbreaks (yellow, light red, dark red taxa) which group in three different clusters that were clearly not related to the 11 infected farms described above (pink taxa).

## Discussion

In this study, we aimed to amplify in a single multiplex PCR reaction a large range of AIV strains. We focused first on those belonging to the H5 subtype. Sequencing the HA was a priority and the *in-silico* design was performed to guarantee, at the minimum, the amplification of the cleavage site of the HA gene for pathotyping. As the viral loads and the integrity of the RNA directly extracted from field samples could be seriously weak or damaged, the PCR panel was designed to produce small (i.e. < 250 bp) to large (>700 bp) amplicons that systematically encompassed the HA cleavage site. Moreover, universal AIV PCR primers were included for a broad-range detection of all type A influenza viruses. While official analyses require tracheal or cloacal swabs, alternative samples have shown promising performances for AIV detection, such as feathers ^23^ and dust sampling in poultry farms ^24^. These samples were included in this study and showed results that were comparable to swabs.

Based on an amplicon-based approach, the workflow described here is intended to focus on influenza viruses with an increased sensitivity, and is not designed for an unbiased metagenomic analysis. This is intentional as AIVs, due to their pathological and epidemiological significance, require a fully dedicated tool that efficiently provides the data needed for surveillance. Due to the genetic evolution of AIVs, a constant update of the PCR panel will be required.

As summarized in previous studies ^25^, two main approaches can be used for viral detection in animals or their environment. PCR (either monoplex or multiplex) remains the mostly widely used as a surveillance or diagnostic tool due to (i) its specificity and low detection threshold, (ii) the inexpensive and simple application of the method and (iii) the robustness and simplicity of its interpretation. However, some limitations have been pointed out, such as nonspecific hybridization, the limited resolution of the information provided, and the difficulty in detecting highly diverse viruses and those for which detailed genetic information (such as pathotype) is crucial. Moreover, PCR cannot handle the detection of minority variants, which can be relevant for viral surveillance.

For these reasons, monitoring protocols increasingly include the use of Next Generation Sequencing (NGS) or third generation sequencing methods that are known to rapidly provide very high-throughput data. Manufacturers in the molecular biology field are now developing tools enabling any laboratory operator to perform NGS in the laboratory or even in the field. However, although these assays are attractive and perform quite well, their running time and cost render them unfeasible for routine clinical applications. Furthermore, online user-friendly bioinformatics tools such as EPI2ME WIMP workflow do not allow the personalized configuration level that can be reached with open source algorithms which also can be run offline. Offline real-time classification pipelines based on open source tools have already been developed ^26^.

Another critical aspect of NGS is the amount of data required for the detection of scarce pathogens whose nucleic acids can be lost among non-viral sequences. For example, a study on Zika virus sequencing showed that 2 million reads were required for the detection of 223 viral reads ^27^. Since the beginning of metagenomics, the scientific community has done everything possible for the development of enrichment techniques, such as host nucleic acids depletion ^28^ and probe capture ^29^ followed by massive deep sequencing. These techniques are interesting for a global and exhaustive description of pathogen communities present in a clinical or environmental sample. Nonetheless, regardless of the enrichment method and sequencing technology, NGS cannot offer detection levels as sensitive as those obtained with PCR. Because PCR offers a fast, cheap and convenient target enrichment, several studies have already shown the advantages of combining PCR with NGS ^27,30,31^.

However, metagenomics should also be considered for its capacity to detect relevant coinfecting pathogens, as well as potential emerging or re-emerging epizootic microorganisms. A shotgun metagenomic workflow for the detection of a wide variety of microbial genomes would require hundreds of megabases per sample. This kind of protocol therefore cannot be routinely applied in veterinary surveillance, but it could be used for field cases of great clinical or epidemiological importance.

ONT technology is highly scalable, from the smallest consumable unit, namely “Flongle”, to high throughput platforms. The Flongle flow cell used in this study allowed a sequential analysis of the avian influenza outbreaks during the course of the epizootics. In addition to enabling low scale sequencing runs, the separate management of each case was an effective way to prevent cross-contamination. Moreover, while the accuracy of sequencing is often identified as a weakness of nanopore technology, it was demonstrated that a sequencing depth cutoff of 10x resulted in a consensus sequence accuracy of 99.95-100%, identical to Illumina sequences [38]. The bioinformatics pipeline described in this manuscript was deliberately based on open-source algorithms that are freely available on dedicated internet platforms. Based on our dataset, we were able to identify AIV HA subtypes, the sequence of the cleavage site and potential clusters of outbreaks. One of the main concerns voiced by government veterinary services is the need to unveil links between outbreaks. These links may suggest direct or indirect farm-to-farm contamination routes. In contrast, when an emerging strain is identified in a region, this may indicate a new introduction, for example from wild birds.

The protocol presented in this study can be carried out for a set of 30 samples in less than 24 hours, starting from the reception of the samples to the production of the consensus sequences. This turnaround time could, however, be reduced by the use of (i) a simplified extraction process, (ii) an automatized PCR amplification and library preparation (eg: voltrax) and a built-in analysis pipeline such as the RAMPART model ^32^.

We propose here a MinION workflow for the rapid typing of avian influenza viruses, evaluated on clinical samples, without any amplification step on cell cultures or embryonated eggs. The objective of this proof-of-concept study was not to perform a comprehensive phylogenetic survey, but rather to demonstrate the possible contribution of this workflow to the real-time management of viral epizootics by unveiling very quickly possible links and specific patterns of transmission. The implementation of this kind of workflow – from the sample to the phylogeny tree - in the framework of official surveillance will obviously need further validation by reference laboratories, but it addresses a substantial need for the surveillance of AIVs and, more broadly, emerging viruses.

## Methods

### Ethics statement

All samples were collected by official veterinarians in farms suspected of being infected with highly pathogenic avian influenza virus, in the framework of the official surveillance of HPAI, as defined by the national veterinary authorities. All these operations were done under the supervision of veterinary authorities and do not fall under the regulation of experiments on animals, as defined in the UE by the Directive 2010/63/EU.

### Sampling and nucleic acids extraction

In a first attempt, four AIV isolates of highly pathogenic (H5N8) and low pathogenicity avian influenza viruses (H5N3, H6N1, H7N9 and H9N2) (Table 3) were included in a validation assay and were processed under the same conditions described below for the clinical samples.

We conducted our study in seven administrative divisions of southwestern and western France, accounting for more than 60% of French duck production. We sampled 30 flocks and collected tracheal swabs, feathers and environmental samples. The sampling information can be found in Supplementary File S1.

All samples were processed in a BSL3 laboratory until lysis in strict compliance with biosafety procedures. Swabs, feather pulps and wipes were placed in 1X PBS and vigorously vortexed for 30 s before viral RNA extraction using the Nucleospin RNA virus kit (Macherey Nagel). Viral RNA was then stored at -80°C until further use.

### RTqPCR screening

RT-qPCR was performed on 2 μL of viral RNA using the Influenza A ID Gene PCR kit (IDVet) for the detection of all type A influenza viruses. The PCR amplifications were run on a LC96 thermocycler (Roche, Basel, Switzerland) with the following parameters: a reverse transcription step of 10 min at 45°C followed by 10 min at 95°C and 40 cycles of 15 s at 95°C coupled with 60 s at 60°C.

### Multiplex-PCR amplification

Hemagglutinin and whole genome amplification were performed using PCR primers pools made of oligonucleotides selected in the literature (Table 1). The PCR amplifications were performed as follows: extracted RNA was reverse transcribed from 10 μL of RNA using a RevertAid First Strand cDNA Synthesis Kit (Thermofisher, Waltham, USA) with 0.5 μM of specific-primer (MBTuni-12 [5’-ACGCGTGATCAGCAAAAGCAGG-3’])^12^. Two PCR assays were combined. First, the variable genomic regions encompassing the CS region were amplified by a primer set designed to anneal the Hemagglutinin belonging to clade 2.3.4.4 (H5HP). Second, the eight genomic segments of influenza A virus were amplified in a single reaction inspired by Zhou and Wentworth with the following universal primers: MBTuni-12 [5’- ACGCGTGATCAGCAAAAGCAGG] and MBTuni-13 [5’-ACGCGTGATCAGTAGAAACAAGG]^12^. The PCR reactions were performed in a final volume of 20 μL using the Phusion™ High–Fidelity DNA Polymerase (ThermoFisher) with 8 μL of primer mix H5HP (composed of 1 μM of each primer) or 0.5 μM of universal primers. The temperature cycle parameters were 98°C for 30 s, followed by 35 cycles (98°C for 10 s, 58°C for 20 s, and 72°C for 3 min) with the final extension at 72°C for 10 min. To assess the PCR amplification products, 5 μL of each PCR product was checked on an agarose gel to roughly estimate the amplicons’ sizes. Each sample produced satisfying amplicons for submission to nanopore sequencing. Such information is afterward critical for the library molarity calculation, which is based on amplicon size and the dsDNA concentration of the PCR products.

### Nanopore sequencing

PCR products were pooled and purified with a 1:1 ratio of AMPure XP beads. DNA libraries were prepared from 200 fmol of purified PCR products using an SQK-LSK109 Ligation sequencing kit supplied by ONT (Oxford Nanopore Technologies, Oxford, United Kingdom) associated with an EXPNBD104 (ONT) native barcoding kit for the multiplexing of samples. For each run, 20 fmol DNA libraries were loaded on a FLO-FLG001 flongle and were run on a MinION Mk1C device (ONT) for 10 hours. High accuracy base-calling was performed in real-time with Guppy (v3.5) embedded in the MK1C software (v19.12.12) with the ‘Trim Barcode’ option on (ONT).

### Sequencing data analysis

The quantity and quality of data were checked by looking at the sequencing report generated by MinKNOW 2.0 and using the NanoPlot from the NanoPack tool set ^33^. The fastq files were aligned using minimap2 ^34^ and the samtools ^35^ package. The HA and whole-genome consensus sequences then were generated using the consensus command of the iVar ^36^ pipeline that embeds the samtools mpileup tool ^37^. The consensus sequences were then manually checked using Bioedit (v7.2.6) and IGV (v2.8.2) software. Nanoplot (Python v3.6.3) and BBmap (v.38.31) were used to assess the quality of the NGS reads and to visualize the coverage of the genome.

The command lines were used as follows:

minimap2 -ax map-ont *reference*.fa *mysample*.fastq > *mysample*.sam

samtools sort *mysample*.sam > *mysample*.sort.bam

samtools index *mysample*.sort.bam

samtools flagstat *mysample*.sort.bam > *mysample*_flagstat

samtools depth *mysample*.sort.bam > *mysample*_samdepth

samtools mpileup -A -Q 0 *mysample*.sort.bam | ivar consensus -p prefix -q 10 -t 0 -m 1

### Phylogeny

Two phylogenetic trees were built with (i) a set of 282 HA sequences from the 2020/21 H5N8 epizootics (including the 11 sequences from this study) and (ii) a set of 91 HA sequences from the 2021/22 H5N1 epizootics (including the 19 sequences from this study). Both sets of sequences are available on the GISAID EpiFlu™ Database and are listed in Supplementary File S6.

The sequences were selected on the basis of their sampling locations and dates and were processed as follows: multiple sequences alignment with MAFFT version 7 ^38,39^, Neighbor-joining phylogenetic tree construction with Jukes-Canto substitution model. Timetrees were then inferred as follows using TreeTime v0.9.5 ^40^:

treetime --aln <input.fasta> --tree <input.nwk> --dates <dates.csv>.

Finally, the tree figures were generated using FigTree v1.4.4.

## Acknowledgements

This study was performed in the framework of the “*Chaire de Biosécurité et Santé Aviaires*”, hosted by the National Veterinary College of Toulouse (ENVT) and funded by the *Direction Generale de l’Alimentation, Ministère de l’Agriculture et de l’Alimentation*, France.

We are grateful to the genotoul bioinformatics platform Toulouse Occitanie (Bioinfo Genotoul, https://doi.org/10.15454/1.5572369328961167E12) for providing help and/or computing and/or storage resources

We would like to thank farmers for their collaboration with the sampling of their flocks.

## Author contributions

G.C. and J.L.G. designed the study; G.C., M.W., L.L., S.S. and F.F. contributed to the experiments and/or their critical analysis; G.C. and JLG wrote the paper. All authors reviewed and approved the manuscript.

## Data availability

The datasets generated during and/or analyzed during the current study are available from the corresponding author on reasonable request.

## Additional information

The authors declare no competing interests.

