## Supplementary File S1 for "An amplicon-based nanopore sequencing workflow for rapid tracking of avian influenza outbreaks, France, 2020-2022"

**Supplementary_file_1_2020-2022 Outbreaks sequencing**

| Reference | Sample type | Sampling date (YYYY-MM-DD) | Host | French department | RTqPCR CT value (M gene) | Total yield (Mb) | N50 (bp) | HA accession number | HA yield (Mb) | N50 HA (bp) |
|---|---|---|---|---|---|---|---|---|---|---|
| 20323 | Tracheal swab | 2020-12-06 | Mule_duck | 40 | 28,5 | 230,59 | 640 | MZ166252 | 30,53 | 661 |
| 20335 | Tracheal swab | 2020-12-10 | Mule_duck | 40 | 24,7 | 139,52 | 891 | MZ166260 | 6,05 | 636 |
| 20338 | Tracheal swab | 2020-12-13 | Mule_duck | 85 | 27,1 | 96,09 | 572 | MZ166268 | 0,36 | 469 |
| 20339 | Dust | 2020-12-13 | Mule_duck | 79 | 20,4 | 129,52 | 882 | MZ166276 | 9,10 | 641 |
| 20347 | Tracheal swab | 2020-12-15 | Mule_duck | 40 | 14 | 474,13 | 937 | MZ166239 | 45,34 | 1 818 |
| 20349 | Tracheal swab | 2020-12-21 | Mule_duck | 40 | 20,4 | 2,34 | 896 | MZ166284 | 0,17 | 656 |
| 20352 | Tracheal swab | 2020-12-26 | Mule_duck | 40 | 18,9 | 289,83 | 808 | MZ166292 | 22,99 | 648 |
| 20353 | Feather | 2020-12-27 | Mule_duck | 40 | 23,5 | 301,95 | 897 | MZ166300 | 28,36 | 598 |
| 21061 | Dust | 2021-01-28 | Mule_duck | 65 | 25 | 28,62 | 443 | MZ166240 | 1,17 | 681 |
| 21064 | Tracheal swab | 2021-01-30 | Mule_duck | 65 | 20 | 28,35 | 872 | MZ166308 | 7,65 | 652 |
| 21084 | Tracheal swab | 2021-02-20 | Mule_duck | 32 | 22,8 | 366,39 | 888 | MZ166316 | 32,68 | 673 |
| 21328 | Dust | 2021-11-28 | Chicken | 59 | 26,23 | 88,81 | 913 | OQ632895 | 81,2 | 915 |
| 21343 | Feather | 2021-12-16 | Mule_duck | 32 | 16,02 | 65,13 | 881 | OQ632818 | 30,24 | 654 |
| 21347 | Feather | 2021-12-18 | Mule_duck | 40 | 16,21 | 153,82 | 713 | OQ632819 | 79,53 | 658 |
| 21348 | Feather | 2021-12-18 | Mule_duck | 64 | 13,03 | 205,89 | 682 | OQ632820 | 121,51 | 611 |
| 21349 | Feather | 2021-12-20 | Mule_duck | 64 | 13,23 | 107,84 | 663 | OQ632821 | 62,74 | 615 |
| 21350 | Dust | 2021-12-23 | Mule_duck | 64 | 21,39 | 76,67 | 905 | OQ632822 | 12,81 | 731 |
| 21351 | Feather | 2021-12-26 | Mule_duck | 40 | 18,14 | 76,03 | 883 | OQ632823 | 33,46 | 646 |
| 21352 | Dust | 2021-12-26 | Mule_duck | 40 | 23,79 | 87,02 | 616 | OQ632824 | 14,78 | 660 |
| 21356 | Feather | 2021-12-30 | Mule_duck | 40 | 17,56 | 99,36 | 890 | OQ632825 | 45,56 | 664 |
| 22005 | Tracheal swab | 2022-01-04 | Pekin_duck | 32 | 17,97 | 83,93 | 903 | OQ632826 | 33,8 | 667 |
| 22008 | Feather | 2022-01-09 | Mule_duck | 40 | 18,76 | 86,05 | 765 | OQ632827 | 40,5 | 661 |
| 22010 | Dust | 2022-01-03 | Mule_duck | 64 | 26,24 | 71,57 | 738 | OQ632828 | 15,66 | 662 |
| 22027 | Feather | 2022-01-11 | Mule_duck | 32 | 18,34 | 98,32 | 887 | OQ632829 | 48,77 | 664 |
| 22029 | Feather | 2022-01-14 | Mule_duck | 64 | 17,12 | 58,08 | 875 | OQ632830 | 29,09 | 648 |
| 22030 | Feather | 2022-01-14 | Mule_duck | 64 | 18,33 | 70,17 | 721 | OQ632831 | 36,26 | 648 |
| 22036 | Feather | 2022-01-19 | Mule_duck | 32 | 20,27 | 72,18 | 738 | OQ632832 | 32,74 | 655 |
| 22077 | Cloacal swab | 2022-02-04 | Pekin_duck | 40 | 27,71 | 78,84 | 630 | OQ632833 | 11,39 | 658 |
| 22083 | Tracheal swab | 2022-02-11 | Mule_duck | 65 | 24,69 | 79,12 | 646 | OQ632834 | 6,6 | 656 |
| 22084 | Feather | 2022-02-11 | Turkey | 85 | 21,02 | 49,45 | 894 | OQ632835 | 18,42 | 652 |
