## Supplementary figures and images for "An amplicon-based nanopore sequencing workflow for rapid tracking of avian influenza outbreaks, France, 2020-2022"

### Supplementary File S2

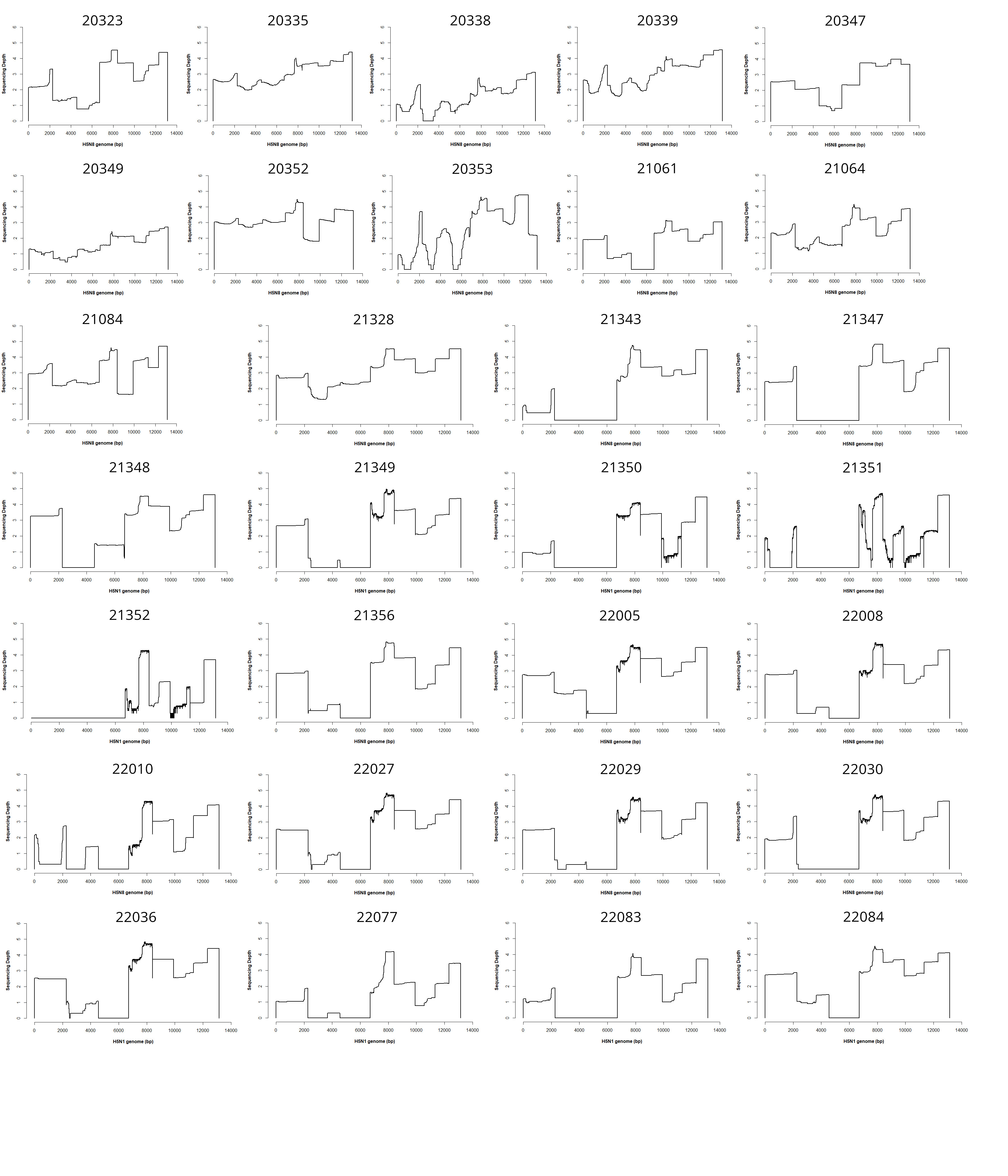

### Supplementary File S4

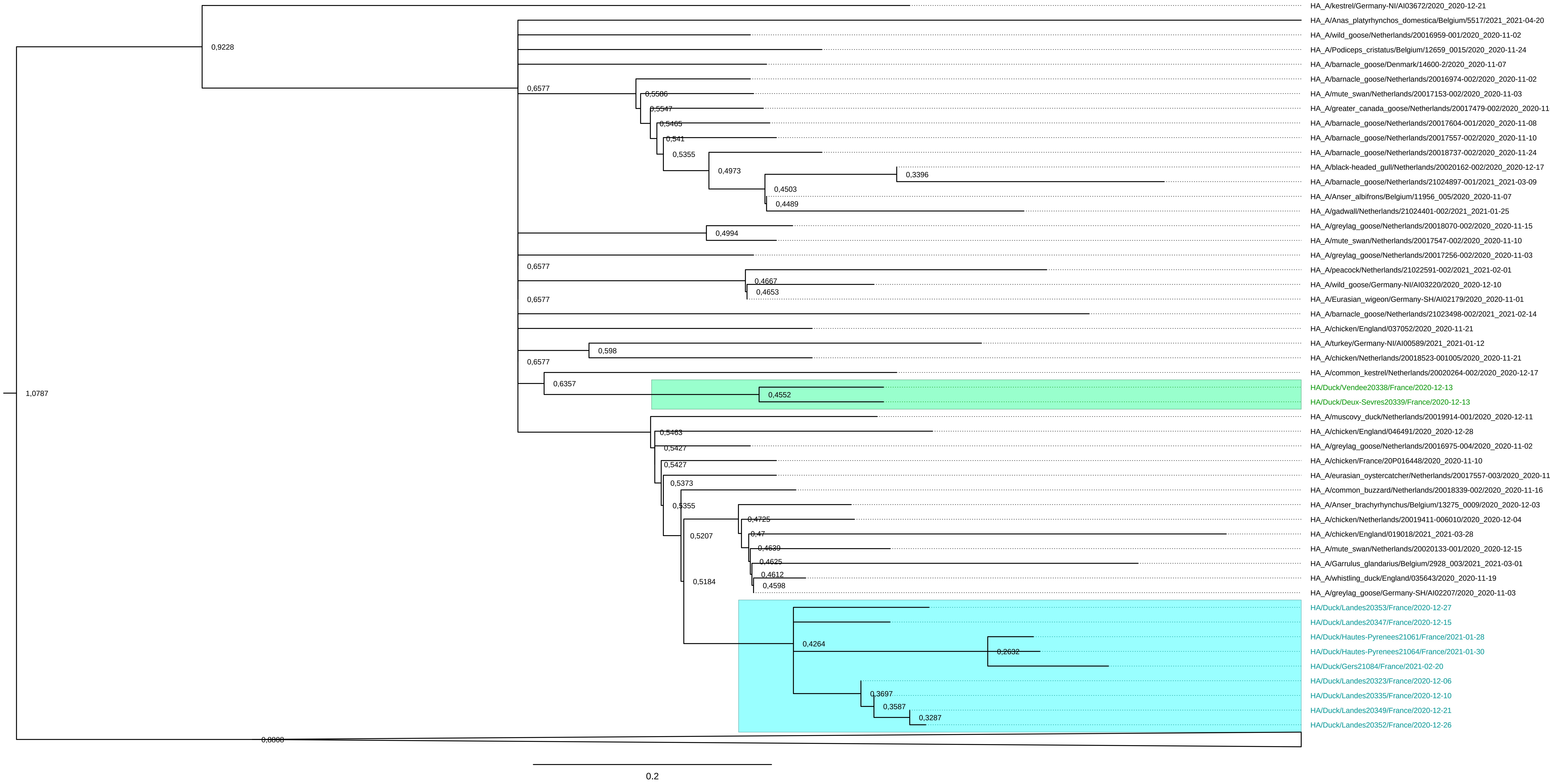

### Supplementary File S5

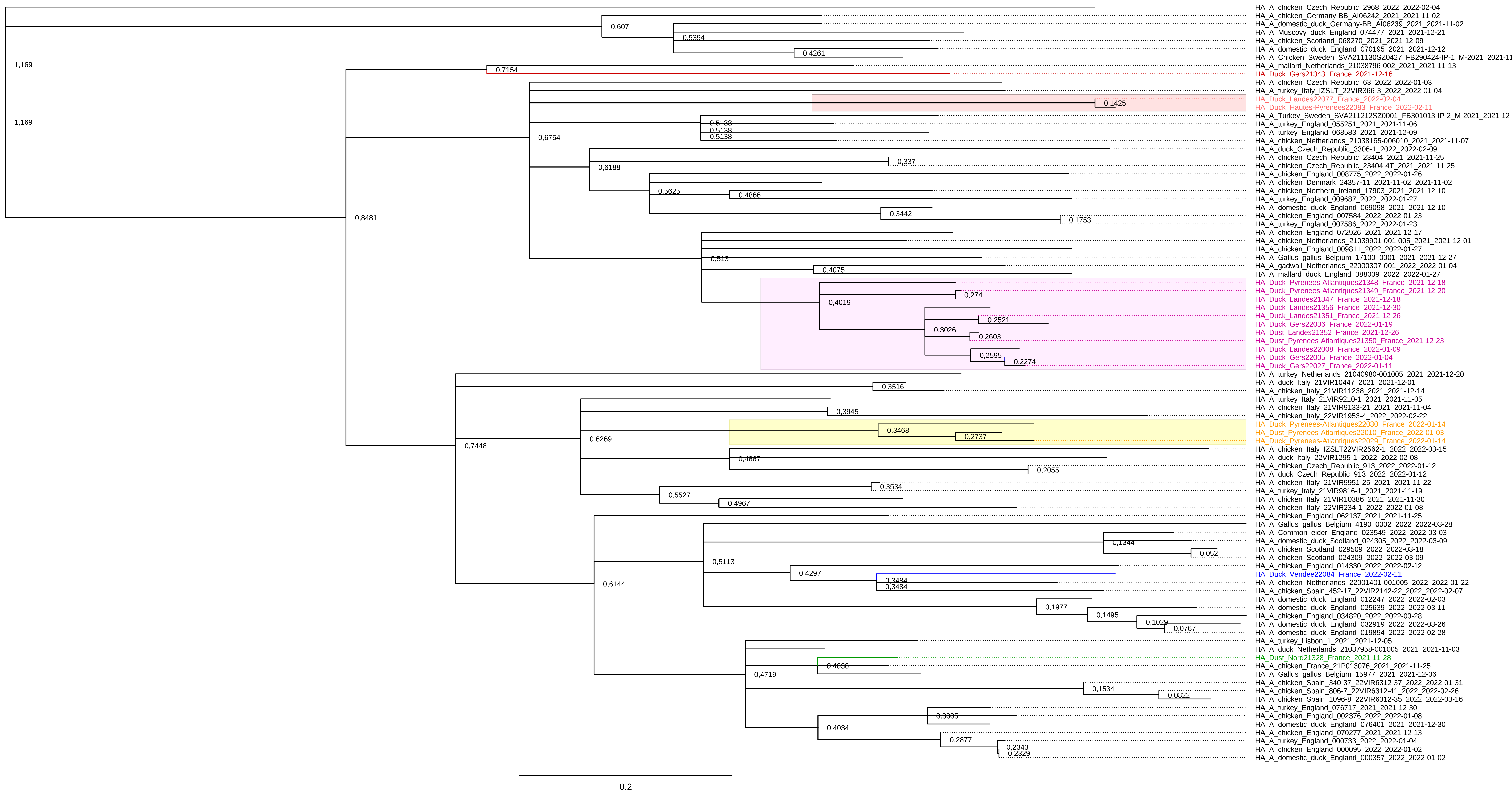
